# Therapeutic potential of inhibition of the neuroinflammation induced cathepsin X: *in vivo* evidence

**DOI:** 10.1101/513671

**Authors:** Anja Pišlar, Larisa Tratnjek, Gordana Glavan, Nace Zidar, Marko Živin, Janko Kos

**Author notes:** These authors contributed equally (Anja Pišlar, Larisa Tratnjek). **Correspondence to:** Anja Pišlar, Faculty of Pharmacy, University of Ljubljana, Aškerčeva 7, 1000 Ljubljana, Slovenia. **Abbreviations:** 6-OHDA, 6-hydroxydopamine; AD, Alzheimer’s disease; BSA, bovine serum albumin; Cc, corpus callosum; CNS, central nervous system; Cpu, striatum; Ctx, cortex; DA, dopamine; DMSO, dimethyl sulfoxide; GFAP, glial fibrillary acidic protein; GPe, external globus pallidus; HRP, horseradish peroxidase; iNOS, inducible nitric oxidase synthase; KPBS, potassium phosphate buffer saline; LPS, lipopolysaccharide; NaPBS, sodium phosphate buffer saline; PD, Parkinson’s disease; SNc, substantia nigra compacta; SVZ, subventricular zone; TH, tyrosine hydroxylase.

## Abstract

Parkinson’s disease (PD) is a progressive neurodegenerative disorder with unknown cause, but it has been postulated that chronic neuroinflammation may play a role in its pathogenesis. Microglia-derived lysosomal cathepsins have been increasingly recognized as important inflammatory mediators. Here, we analyzed the regional distribution and cellular localization of the cathepsin X in the rat brain with neuroinflammation-induced neurodegeneration. Unilateral injection of lipopolysaccharide (LPS) into the striatum induced strong upregulation of cathepsin X expression and its activity in the ipsilateral striatum. In addition to the striatum, cathepsin X overexpression was detected in other areas such as cerebral cortex, corpus callosum, subventricular zone and external globus pallidus mainly restricted to glial cells. Moreover, continuous administration of the cathepsin X specific inhibitor AMS36 showed protective effects against LPS-induced striatal degeneration, as seen by the decreased extent of striatal lesion and decreased expression of neuroinflammation marker. These results demonstrate that glial upregulated cathepsin X may play a role as a potential pathogenic factor in PD. Inhibition of cathepsin X enzymatic activity thus may be useful in preventing neuroinflammation-induced neurodegeneration.

## Introduction

Parkinson’s disease (PD) is a common neurodegenerative disorder characterized by progressive degeneration of the dopaminergic projection between the substantia nigra *compacta* (SNc) and the striatum and by a decrease in striatal dopamine (DA), which is associated with motor impairments (Braak et al, 2003; Hornykiewicz, 1993; Youdim & Riederer, 1997). Although the etiology of PD is not fully understood, accumulating evidence suggests that neuroinflammatory processes, which are mainly mediated by activated microglia and astrocytes, are crucial for the initiation and progression of PD (McGeer et al, 2005; McGeer & McGeer, 2008). During neuroinflammation, activated microglia and astrocytes release a variety of cytokines, chemokines and toxic factors, all of which may lead to subsequent neuronal toxicity and aggressive neuronal loss that drives the pathogenic progress of PD (Choi et al, 2009; Menza et al, 2010; More et al, 2013; Niranjan, 2014). Therefore, the characterization of the endogenous biomolecules involved in neuroinflammation-induced neurodegeneration may be critical in the development of novel therapeutic strategies for treatment of PD.

Increasing evidence shows that activated microglia induce the synthesis and secretion of lysosomal cathepsins, in particular during the early stage of neuroinflammation, to trigger signaling cascades that aggravate neurodegeneration (Clark & Malcangio, 2012; Fan et al, 2015; Fan et al, 2012; Hafner et al, 2013; Kingham & Pocock, 2001; Pislar et al, 2017; Terada et al, 2010; Wendt et al, 2009). To date, most neuroinflammation research has focused on cysteine cathepsins, which are the largest cathepsin family, comprising 11 members (cathepsins B, C, F, H, K, L, O, S, W, V and X). These cathepsins possess a conserved active site involving cysteine, histidine and aspargine residues in a catalytic triad (Rawlings et al, 2012). They are primarily responsible for terminal protein degradation in lysosomes; however, their role in regulating a number of other important physiological and pathological processes has been demonstrated (Obermajer et al, 2006; Pislar & Kos, 2014; Pislar et al, 2015; Stoka et al, 2016).

The role of cathepsin X, a cysteine cathepsin with solely carboxypeptidase activity, has been chiefly restricted to cells of the immune system, predominantly monocytes, macrophages and dendritic cells (Kos et al, 2009; Kos et al, 2005). Cathepsin X expression and proteolytic activity have also been found to be strongly upregulated in mouse brain, with a preference for glial cells and aged neurons (Stichel & Luebbert, 2007; Wendt et al, 2007). In a transgenic mouse model of Alzheimer’s disease (AD), cathepsin X upregulation in microglial cells surrounding amyloid plaques has been observed, and further, the involvement of cathepsin X in amyloid-β-related neurodegeneration through proteolytic cleavage of the C-terminal end of γ-enolase has been suggested (Hafner et al, 2013; Pislar & Kos, 2013; Wendt et al, 2007). Additionally, cathepsin X also promotes the apoptosis of neuronal cells induced by the neurotoxin 6-hydroxydopamine (6-OHDA), which was reversed with cathepsin X downregulation or inhibition by the specific cathepsin X inhibitor (Pislar et al, 2014). Moreover, *in vivo* cathepsin X expression and its activity have been found to be strongly increased in the SNc of hemi-parkinsonian rats with 6-OHDA-induced excitotoxicity in the unilateral medial forebrain bundle, indicating that cathepsin X may be involved in the pathogenic cascade event in PD (Pislar et al, 2018). In addition to a role in neurodegeneration, the involvement of cathepsin X in inflammation-induced neurodegeneration has been reported. Substantially increased secretion of cathepsin X from microglia has been observed in response to the inflammatory stimulus induced by lipopolysaccharide (LPS), leading to microglia activation-mediated neurodegeneration (Pislar et al, 2017; Wendt et al, 2009). To date, the cathepsin X expression and its role in inflammation-induced neurodegeneration have been studied mainly *in vitro*; however, the expression pattern and cellular localization of cathepsin X in the brain, as well as the functional role of cathepsin X in chronicle inflammation related to neurodegeneration still remain unclear.

The current study explored the expression pattern and cellular localization of cysteine protease cathepsin X in the rat brain following LPS-induced striatal inflammation. Cathepsin X was found to be highly overexpressed after LPS injection and its upregulation was restricted to microglia cells and reactive astrocytes in the ipsilateral striatum and distributed throughout the different areas, including the cortex, corpus callosum and external globus pallidus. To gain further insight into the role of upregulated cathepsin X in neuroinflammation-induced neurodegeneration, rats were treated with cathepsin X-specific inhibitor AMS36, which proved to be partially effective in protecting against LPS-induced neuroinflammation.

## Results

### Neuroinflammation-induced cathepsin X expression in the brain

Rats were lesioned by an injection of LPS into the right striatum to evaluate cathepsin X expression pattern in the brain with the inflammation-induced neurodegenerative pathology. Firstly, assessment of striatal degeneration showed a significant reduction in striatal size and increased lateral ventricle size on the ipsilateral side following LPS injection (Suppl. Fig 1). Immunohistochemical analysis of TH expression, a dopaminergic neuronal marker, revealed an abundant loss of TH-positive fibers in the ipsilateral striatum after LPS injection (Fig 1A), while no loss of TH-positive cells in the ipsilateral SNc was observed as compared to the contralateral side (Fig 1B). In contrast, the results of immunohistochemical analysis of cathepsin X showed an increase of protein expression in the ipsilateral striatum and surrounding areas (Fig 1A’), whereas no increase in cathepsin X immunoreactivity was observed in the ipsilateral SNc four weeks after LPS injection (Fig 1B’). Likewise, immunofluorescence staining of the ipsilateral striatum showed decreased and patchy TH-immunoreactivity, with a concomitant marked increase of cathepsin X expression (Fig 1C). Immunofluorescencent staining also showed no upregulation of cathepsin X expression in the ipsilateral SNc, which coincides with no change in TH-immunostaining intensity in the ipsilateral SNc when compared to the contralateral side (Fig 1D).

**Figure 1.**
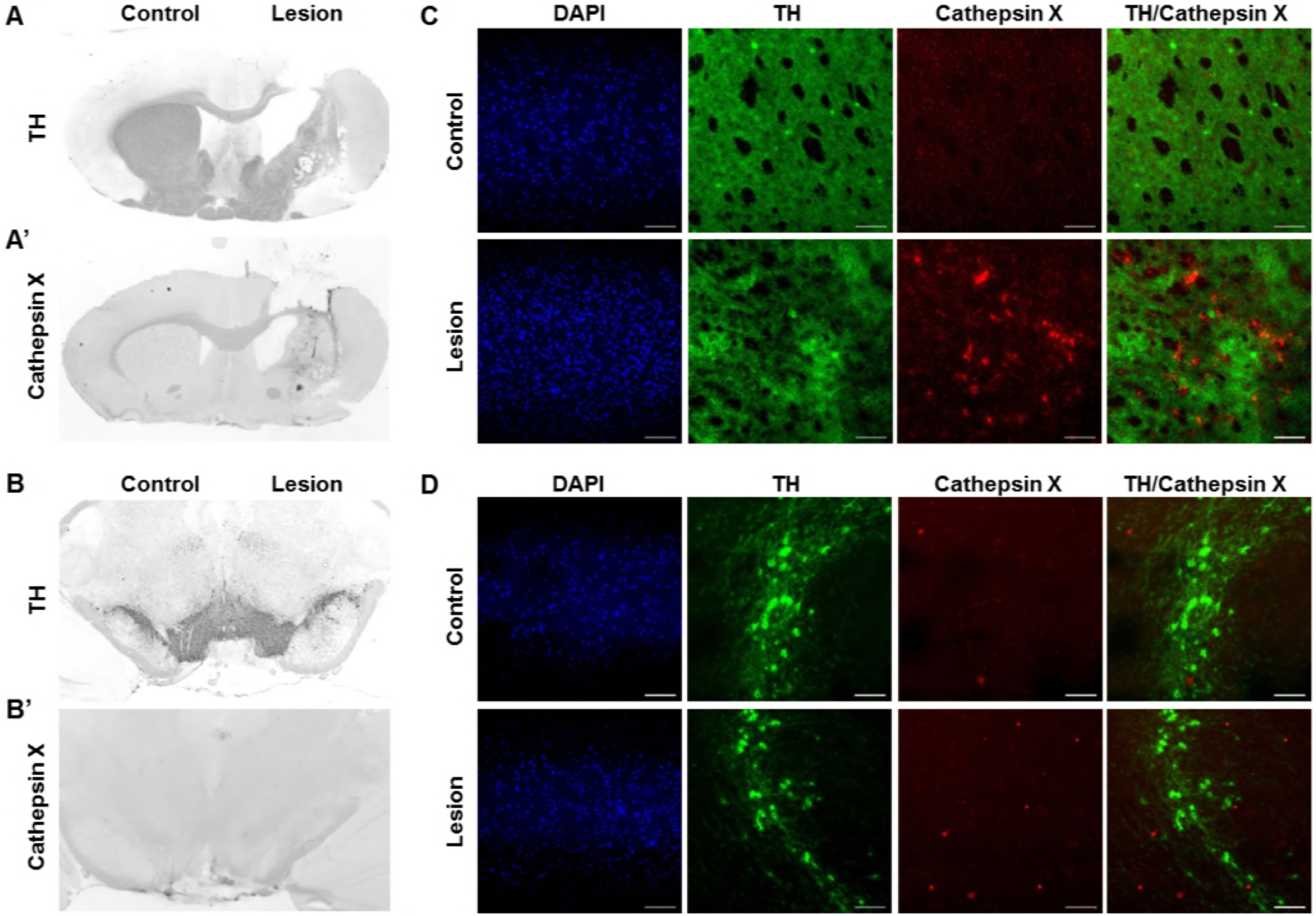
Cathepsin X upregulation in the striatum after intrastriatal LPS injection. (A, B) Representative immunohistochemical images of coronal sections of the striatum (A) and SNc (B) from the four weeks’ time-point after LPS-induced lesion. TH staining demonstrated decrease of TH-positive fibers in the striatum (A) and no loss of TH-positive cells in the SNc (B, *upper panel*) in the ipsilateral side (Lesion) compared to contralateral side (Control), whereas cathepsin X immunoreactivity was markedly increased in the ipsilateral striatum after LPS injection (A’). No increase in cathepsin X immunoreactivity was observed in the ipsilateral SNc after LPS-induced lesion (B’, *lower panel*). (C, D) Representative images of double immunofluorescence staining of TH (green fluorescence) and cathepsin X (red fluorescence) in the striatum (C) and SNc (D) at four weeks after the LPS injection. Nuclei were counterstained with DAPI (blue fluorescence). In the ipsilateral (Lesion) striatum, TH-positive fibers were reduced compared to the contralateral side (Control), whereas cathepsin X expression was upregulated in the ipsilateral side of striatum after LPS injection (C). In the SNc, no upregulation of cathepsin X in the ipsilateral side was observed (D). Group of 5 animals was conducted (n = 5), where 4 anterior-posterior striatal slices and 3 nigral anterior-posterior slices of each animal were analyzed. *Scale bar = 20* μm.

Based on the immunostaining observations of cathepsin X upregulation after LPS-induced lesion, proteins were isolated from the dissected contralateral and ipsilateral striatal and SNc regions of coronal sections in order to confirm changes in cathepsin X protein level and its enzymatic activity using quantitative methods. Western blot analysis revealed a prominent increased expression of cathepsin X protein level in the ipsilateral striatum compared to the contralateral striatum, with the predominant proform of cathepsin X. No significant differences in the expression of cathepsin X were seen between the ipsilateral and contralateral SNc (Fig 2A). ELISA results also confirmed significantly upregulated cathepsin X protein level in the ipsilateral striatum compared to the contralateral side four weeks after LPS injection as shown in Fig 2B. Furthermore, analysis of protein activity showed that the LPS-induced lesion also significantly increased enzymatic activity of cathepsin X in the ipsilateral striatum relative to contralateral (Fig 2C). In contrast, no significant increase in cathepsin X activity was detectable in the ipsilateral SNc compared with the enzyme activity in the contralateral SNc (Fig 2C), which is consistent with the results of cathepsin X protein level in the SNc obtained by both immunostaining and western blot analysis. Taken together, these observations suggest that the LPS-induced striatal degeneration might be associated with an increase in cathepsin X expression and its activity in brain with parkinsonian pathology.

**Figure 2.**
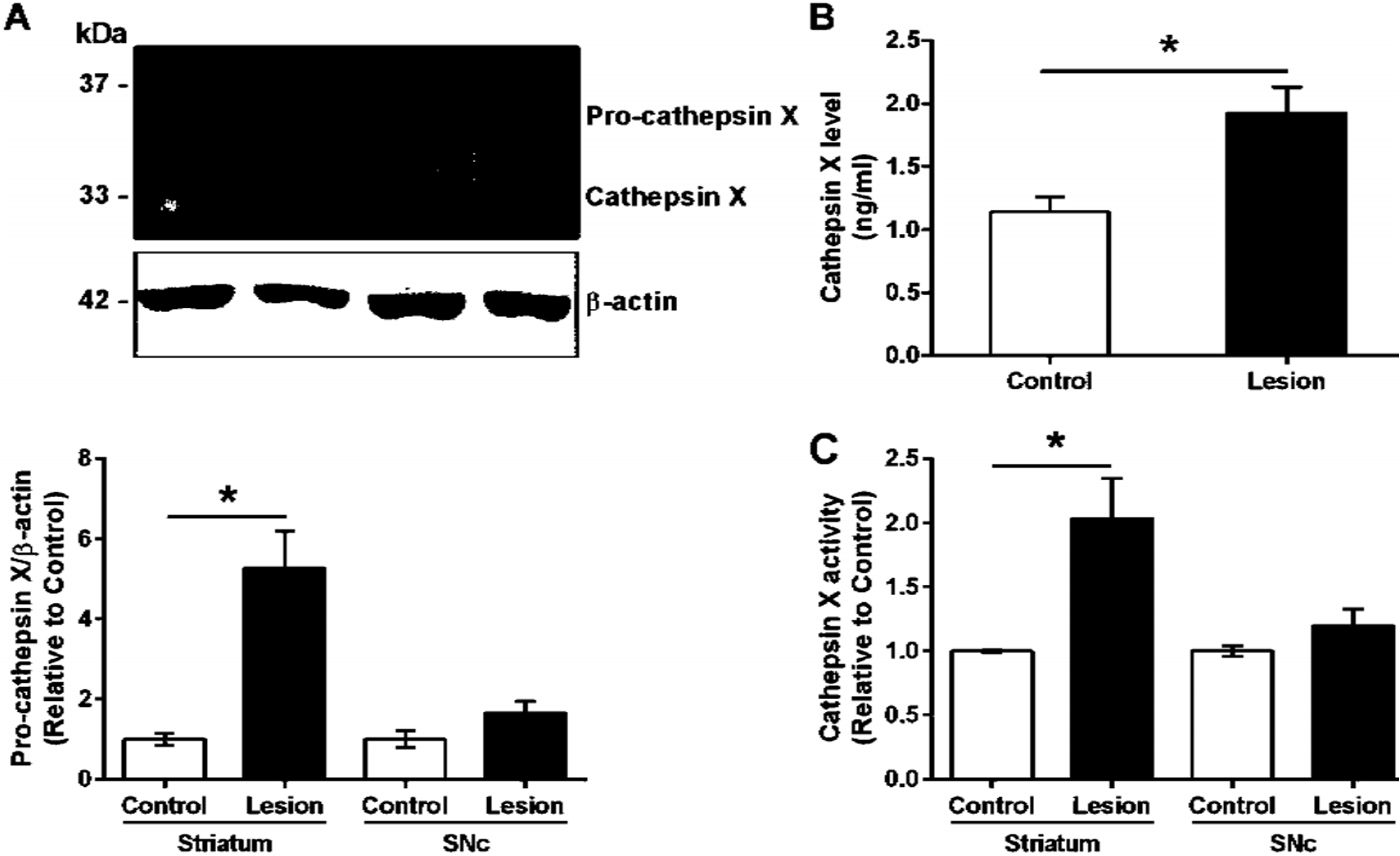
Cathepsin X protein level and activity in the striatum and SNc following intrastriatal LPS injection. Four weeks after the intrastriatal induced LPS lesion, the contralateral (Control) and ipsilateral (Lesion) sides of the striatum and SNc were dissected from coronal sections, homogenized, and analyzed for cathepsin X protein levels and activity. (A) Western blot was performed using goat polyclonal anti-cathepsin X antibody. Rabbit monoclonal antibody raised against β-actin was used as loading control. The graph below blots shows a semi-quantitative densitometry analysis of the protein level relative to that in the contralateral side. Cathepsin X protein level was significantly increased in the ipsilateral striatum compared to contralateral, whereas no significant difference in protein level in the SNc was observed. (B) ELISA analysis of the cathepsin X protein level in the striatum after LPS-induced lesion. Cathepsin X protein level in the ipsilateral striatum was significantly increased compared to the contralateral side. (C) Enzymatic activity analysis shows an increase in cathepsin X activity in the lesioned striatum after LPS injection, whereas the increase in cathepsin X activity in the ipsilateral SNc was not significant compared to the activity in contralateral SNc. Values are means ± SD of group of 5 animals (n = 5), where 4 anterior-posterior striatal slices and 8 nigral anterior-posterior slices of each animal were analyzed (two-tailed Student’s *t-*test, * *p* < 0.05 *vs* Control).

### Distribution of brain cathepsin X following LPS injection

In order to more precisely define the distribution of upregulated cathepsin X in LPS-induced neuroinflammation, we performed additional immunohistochemical staining of cathepsin X on the striatal brain cryosections. No noticeable staining for cathepsin X was detected in the contralateral striatum or throughout the different areas such as cortex (Ctx), corpus callosum (Cc), and external globus pallidus (GPe) (Fig 3A and 3B). However, an abundant increase in cathepsin X expression was observed predominantly in the ipsilateral striatum (Fig 3C, D and G), where condensed expression of cathepsin X was apparent (Fig 3C’) or scattered throughout the striatum region (Fig 3D’, 3G’). Strong upregulation of cathepsin X expression was also observed in the ipsilateral Ctx (Fig 3E) and Cc (Fig 3F) around the injection track, where dense cathepsin X staining was observed. Nevertheless, a marked increase in cathepsin X expression was also observed in the ipsilateral GPe (Fig 3H) with scattered distribution through the region. Thus, results of altered cathepsin X expression and distribution following LPS injection indicate relevant biological function of the protein in the brain under pathological conditions.

**Figure 3.**
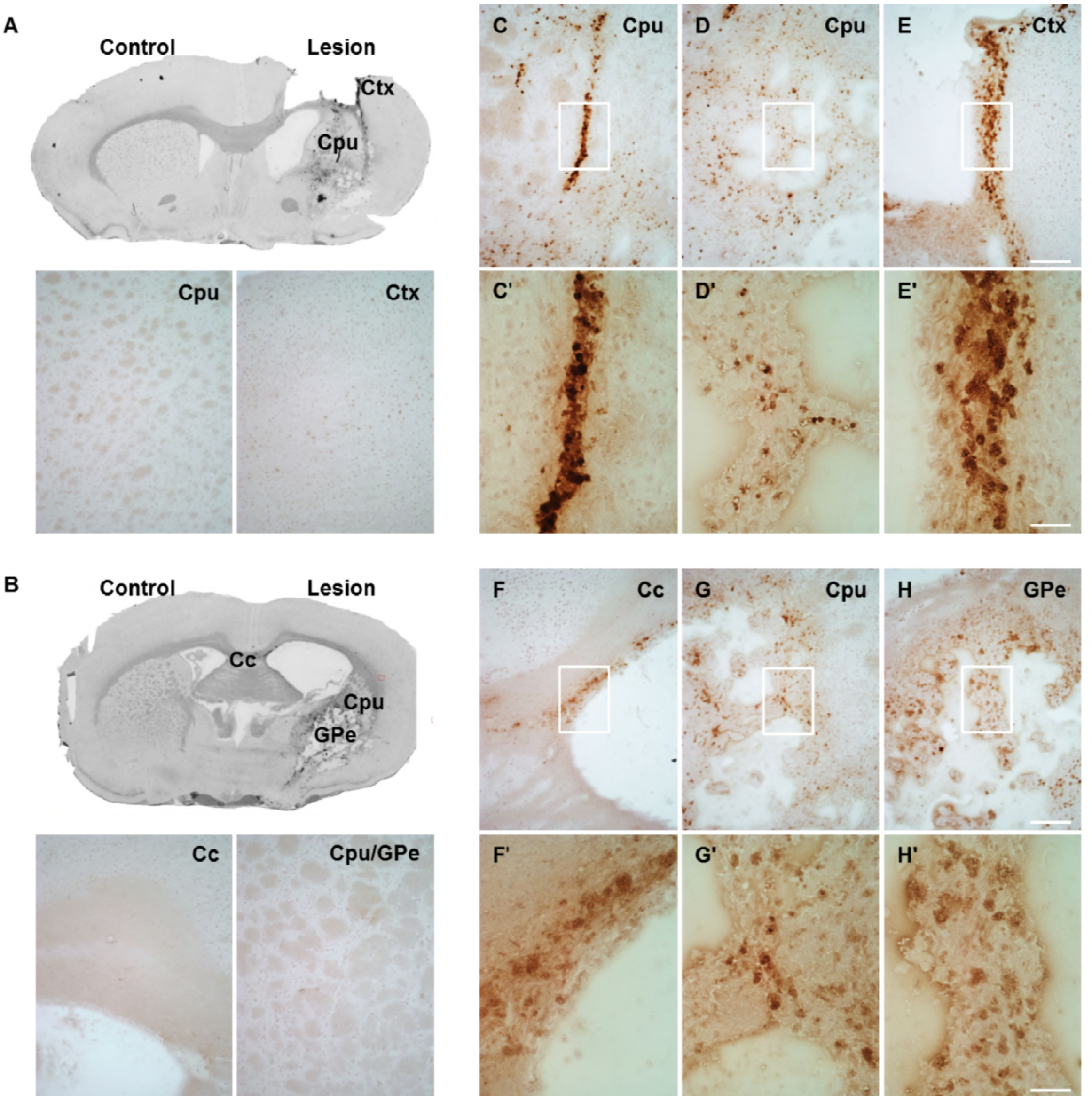
The expression pattern of upregulated cathepsin X after LPS injection. Representative cathepsin X immunohistochemical images of the contralateral (Control) and ipsilateral (Lesion) sides of the striatal brain slices four weeks after LPS-induced lesion. (A, B) In the control, no upregulation of cathepsin X in the striatum (caudate-putamen, Cpu) or surrounding areas was observed. (C-H) Cathepsin X staining demonstrated that cathepsin X expression was strongly induced in the ipsilateral striatum (Cpu; C, D, G) and distributed throughout the different areas, such as the ipsilateral cortex (Ctx; E), corpus callosum (Cc; F) and external globus pallidus (GPe; H). The white box areas in images C-H are shown in magnified view in images C’ - H’. Group of 5 animals (n = 5) was conducted, where 4 anterior-posterior striatal slices of each animal were analyzed. *Scale bar* = 200 μm (C-H); 50 μm (C’-H’).

### Cellular localization of brain cathepsin X following LPS injection

In order to further explore which type of cells express cathepsin X following LPS injection, we conducted double immunofluorescent staining to detect the co-localization of cathepsin X with different cell-specific markers, including markers for neurons (NeuN), microglial cells (CD11b) and astrocytes (GFAP). Weakly expressed cathepsin X in the contralateral striatum was mainly present in NeuN-positive cells (Suppl. Fig 2). However, as shown in Fig 4, four weeks after LPS injection cathepsin X was not upregulated in NeuN-positive cells in the ipsilateral striatum as well as in other areas such as Ctx, Cc, GPe and subventricular zone (SVZ).

**Figure 4.**
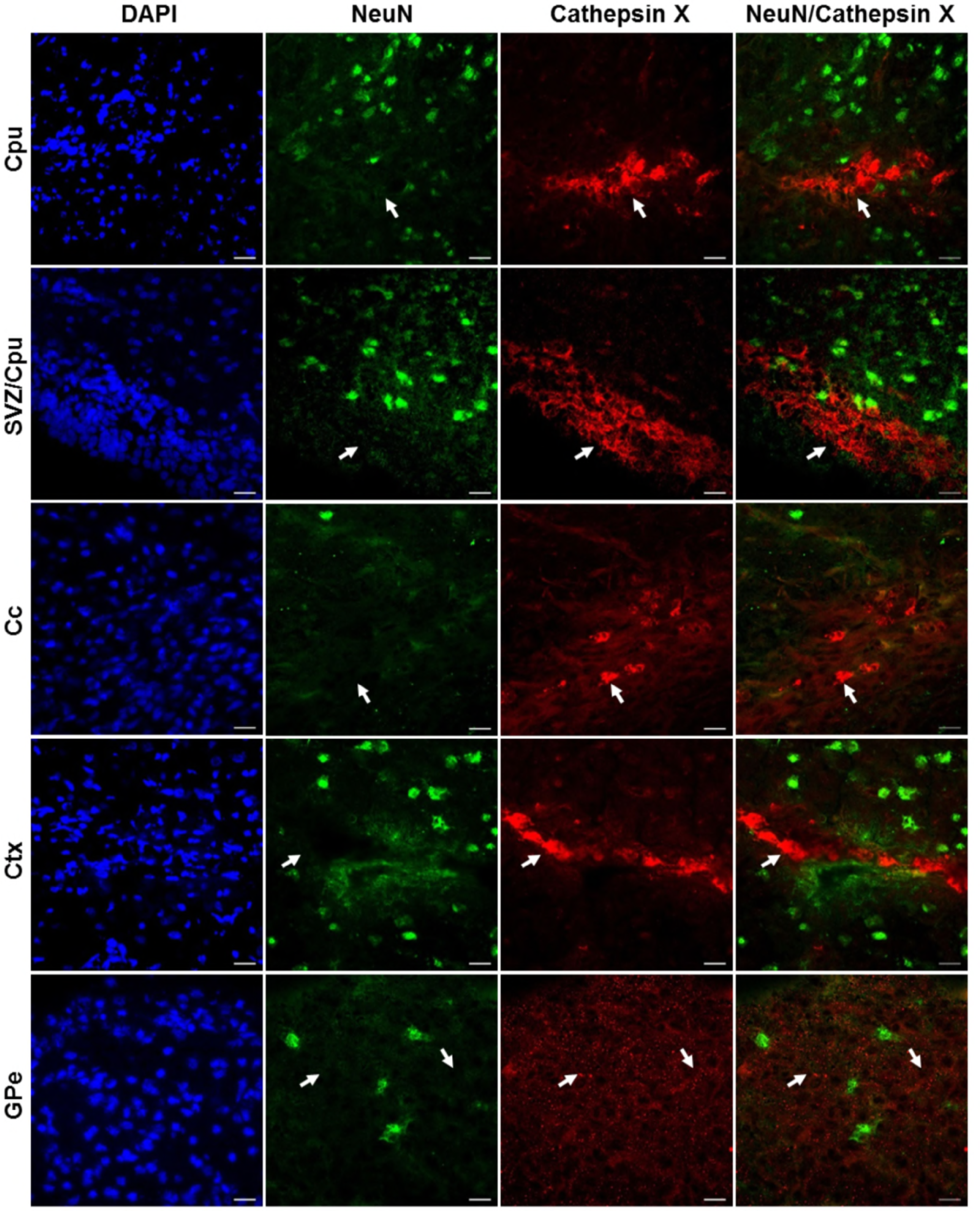
Cathepsin X is not upregulated in neuronal cells after intrastriatal LPS injection. Representative images of double immunofluorescence staining of neuronal marker NeuN (green fluorescence) and cathepsin X (red fluorescence) in the ipsilateral side of the striatal brain slices four weeks after LPS-induced lesion. In the striatum (caudate-putamen, Cpu), subventricular zone (SVZ), cortex (Ctx), corpus callosum (Cc) and external globus pallidus (GPe) cathepsin X-positive signal was strongly upregulated, however, no neuronal cells were positive for cathepsin X (*arrows*). Group of 5 animals (n = 5) was conducted, where 4 anterior-posterior striatal slices of each animal were analyzed. *Scale bar* = 20 μm.

Upregulated cathepsin X on the ipsilateral side after LPS-induced lesion was restricted to glial cells. Cathepsin X staining was localized mainly within microglia cells in the striatum and throughout different areas including Ctx, Cc, GPe and SVZ as demonstrated by co-localization of cathepsin X with CD11b (Fig 5). Strong co-localization of cathepsin X with GFAP was also observed in the SVZ at the ipsilateral side, and some astrocytes in the ipsilateral striatum, Ctx, Cc and GPe also showed cathepsin X overexpression (Fig 6). Nevertheless, after four weeks of LPS-induced lesion, no localization of physiologically expressed cathepsin X was observed in glial cells in the contralateral striatum probably due to minimal glia reactivity, where only individual CD11b- and GFAP-positive cells were observed (Suppl. Fig 2). These results further suggest the significant role of cathepsin X in neurodegeneration that is associated with glial activation.

**Figure 5.**
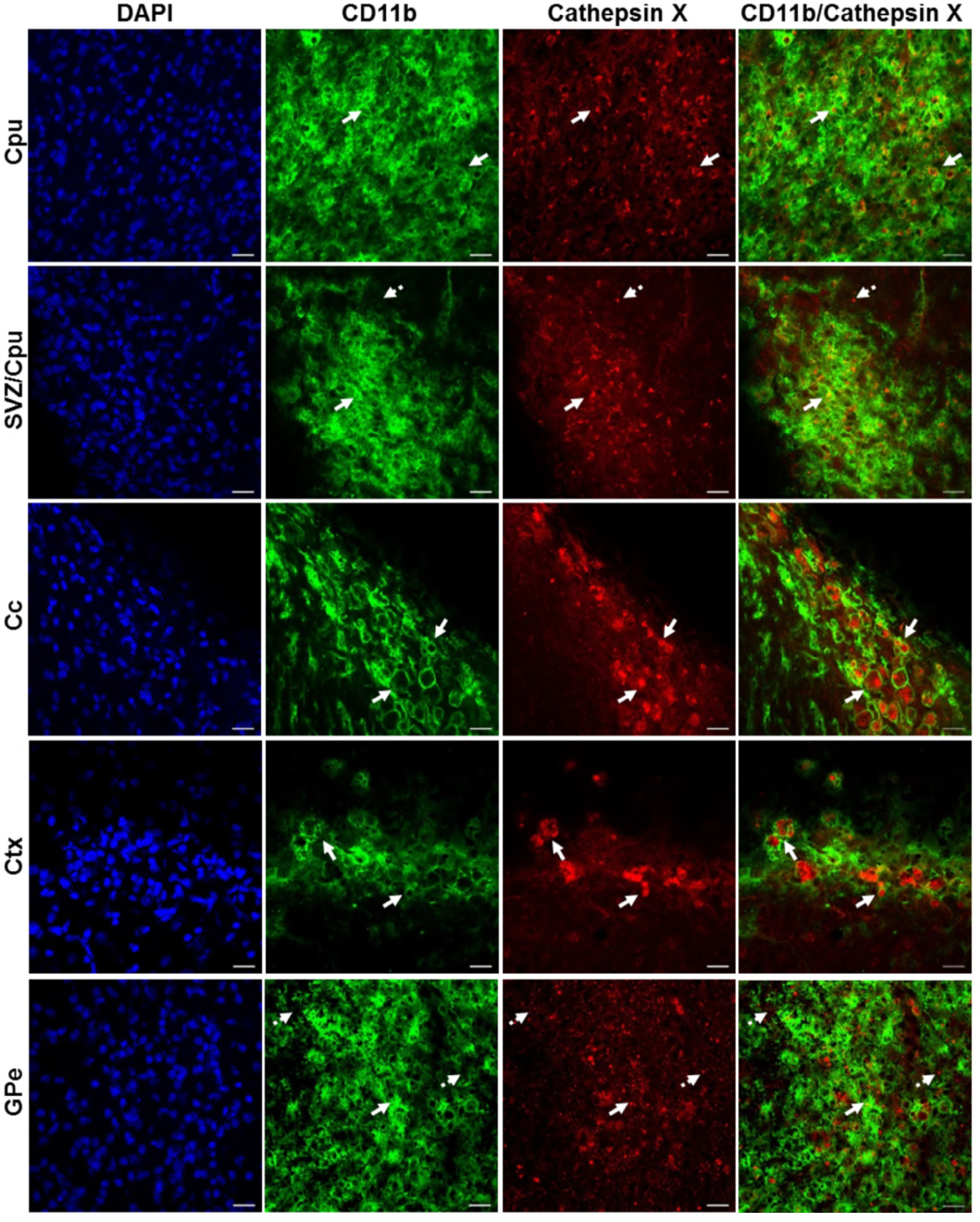
Microglial phenotype of increased cathepsin X-immunopositive cells after intrastriatal LPS injection. Representative images of double immunofluorescence staining of microglial marker CD11b (green fluorescence) and cathepsin X (red fluorescence) in the ipsilateral side of the striatal brain slices four weeks after LPS-induced lesion. In the striatum (caudate-putamen, Cpu), subventriclular zone (SVZ), cortex (Ctx), corpus callosum (Cc) and external globus pallidus (GPe) upregulated cathepsin X was predominantly restricted to CD11b-positive cells (*arrows*) however, some cathepsin X-positive signal did not overlap with CD11b-immunosignal in SVZ/Cpu and GPe (*dashed arrows*). Group of 5 animals (n = 5) was conducted, where 4 anterior-posterior striatal slices of each animal were analyzed. *Scale bar* = 20 μm.

**Figure 6.**
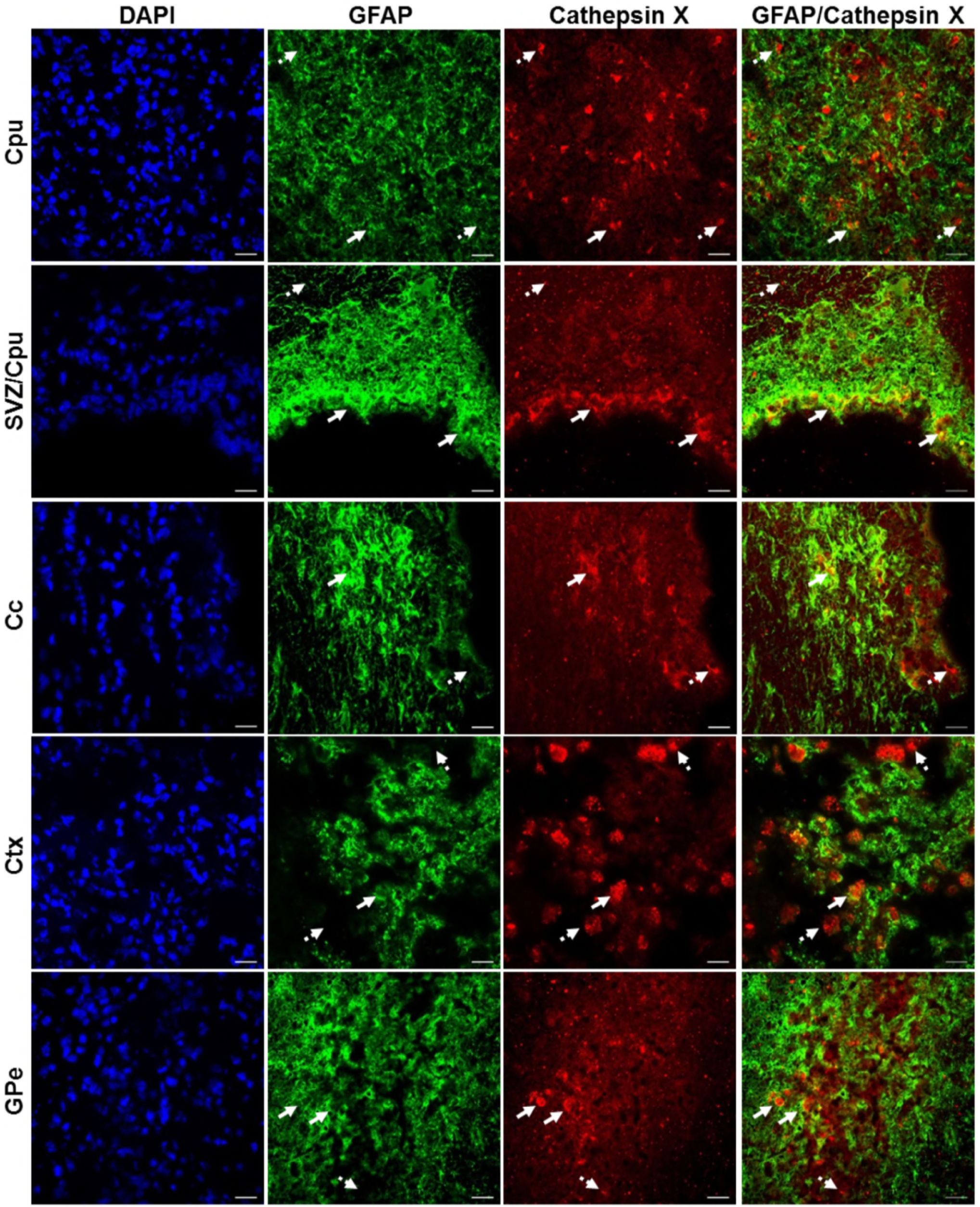
Astroglial phenotype of increased cathepsin X-immunopositive cells after intrastriatal LPS injection. Representative images of double immunofluorescence staining of marker for astrocytes GFAP (green fluorescence) and cathepsin X (red fluorescence) in the ipsilateral side of striatal brain slices four weeks after LPS-induced lesion. In the striatum (caudate-putamen, Cpu), subventricular zone (SVZ) cortex (Ctx), corpus callosum (Cc) and external globus pallidus (GPe) increased cathepsin X expression was partially observed in GFAP-positive cells, predominantly in SVZ/Cpu area (*arrows*); however in all regions cathepsin-positive signal was also seen in GFAP-negative cells (*dashed arrows*). Group of 5 animals (n = 5) was conducted, where 4 anterior-posterior striatal slices of each animal were analyzed. *Scale bar* = 20 μm.

### Evaluation of *in vivo* potency of cathepsin X inhibitor

Intrigued by the results of strongly increased cathepsin X expression and its activity after LPS-induced lesion, we were interested whether the inhibition of cathepsin X enzymatic activity might be useful in preventing the striatal degeneration. First, we determined the potency of cathepsin X inhibition by specific inhibitor AMS36 *in vivo*. After the inhibitor administration, decreased cathepsin X activity was observed in cerebellum extracts; however, a significant decrease in cathepsin X activity was detectable only after 2 days of administration (Suppl. Fig 3). Therefore, to determine the potential effect of AMS36 on LPS-induced striatal degeneration, AMS36 was administrated 2 days prior to LPS injection and then every 4 days within four weeks. As demonstrated by striatal brain sections, AMS36 attenuated the LPS-mediated increase of the lateral ventricles (Fig 7A). As shown in Fig 7B, statistical analysis of the lateral ventricle on the ipsilateral side revealed trend for the reduced size of lateral ventricle in presence of AMS36; however, the difference was not significant. Additionally, there was a trend toward decreased extent of striatal lesion when AMS36 was administrated following LPS injection compared to LPS injection alone (Fig 7C).

**Figure 7.**
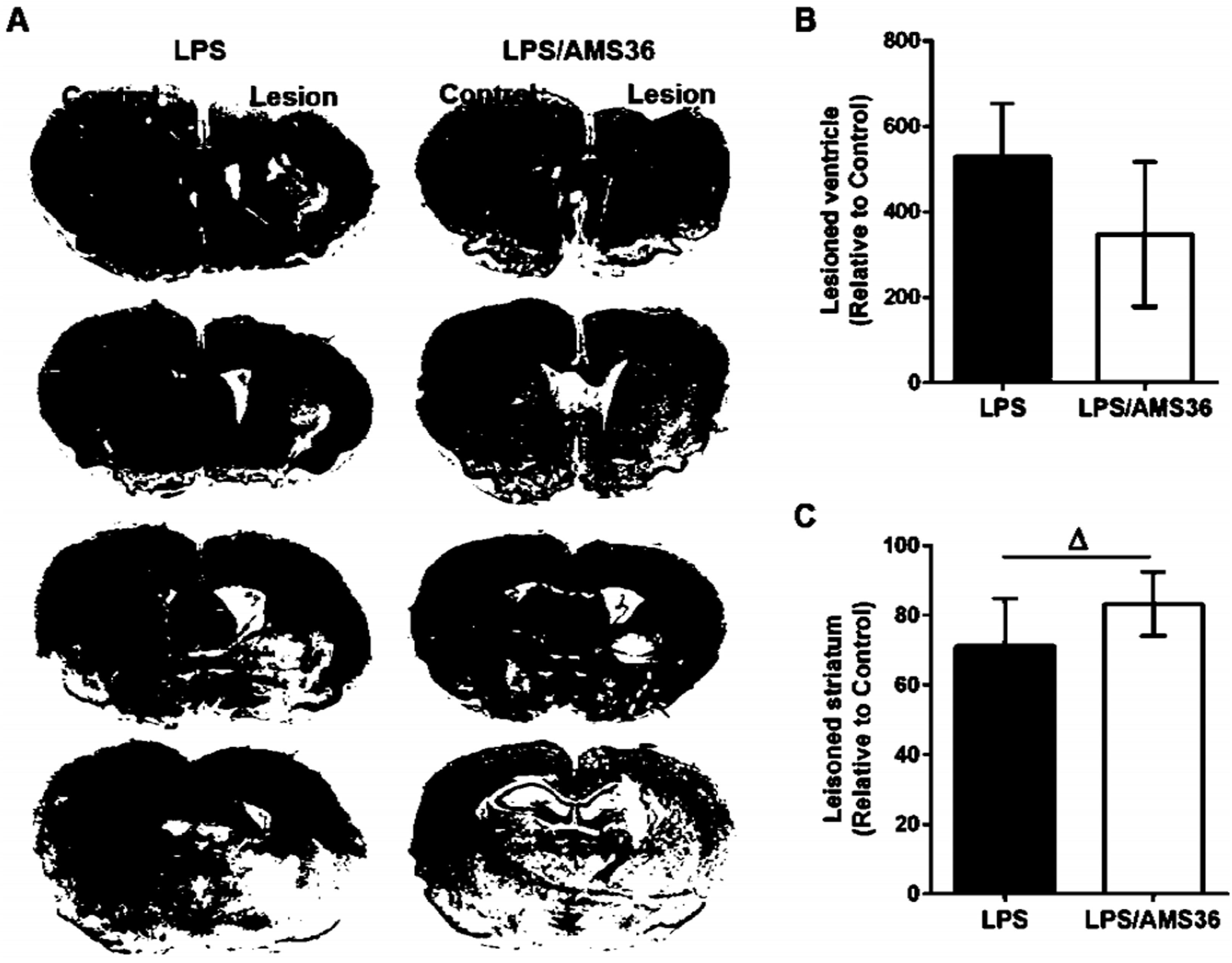
Cathepsin X inhibitor alleviated degeneration following intrastriatal LPS injection. (A) Representative images of methylene blue staining of coronal sections of the striatum cut at anterior-posterior distances four weeks after LPS-induced lesion in the absence of cathepsin X inhibitor AMS36 (LPS) or the presence of AMS36 (LPS/AMS36). (B) The graph shows the analysis of the size of ventricle as a sum of lateral ventricle areas in 4 anterior-posterior striatal slices at the ipsilateral side (Lesion) relative to contralateral side (Control). (C) The graph shows the analysis of the size of striatum as a sum of striatum areas in 4 anterior-posterior striatal slices at the ipsilateral side relative to contralateral side. Values are means ± SD of group of 5 animals for LPS and group of 6 animals for LPS/AMS36, where 4 anterior-posterior striatal slices from each animal were analyzed (n = 5/6) (Bonferroni multiple comparisons *t*-test, ^Δ^ *p* < 0.1 LPS/AMS36 *vs* LPS).

Furthermore, we characterized neuroinflammation by immunostaining of CD11b, a marker for activated microglia, and we observed strong co-localization of cathepsin X with CD11b at the ipsilateral striatum and throughout other areas. iNOS is expressed in microglia during neuroinflammation and produces excessive amount of nitric oxide, which can cause death of dopaminergic neurons (Bal-Price & Brown, 2001). Therefore, the expression patterns of iNOS and TH were examined in the ipsilateral striatum following intrastriatal LPS injection, in the presence or absence of inhibitor AMS36 after four weeks administration. Increased iNOS expression was observed in the ipsilateral striatum following LPS injection compared to contralateral striatum as demonstrated by western blot analysis. The increased striatal iNOS returned to control level when AMS36 was continuously administrated together with LPS injection. However, unfortunately there was no statistically significant difference in iNOS expression in the ipsilateral striatum following LPS/AMS36 treatment compared to protein level of iNOS in the ipsilateral striatum following LPS injection alone (Fig 8A). Moreover, cathepsin X inhibitor also affected TH protein level after LPS-induced lesion. LPS injection for four weeks caused significant decrease in TH protein level in the ipsilateral striatum compared to contralateral, whereas TH protein expression was not significantly reduced in the ipsilateral striatum when AMS36 was administrated (Fig 8B). It was therefore demonstrated that cathepsin X inhibitor shows a potential protective effect against striatal degeneration, which also supports a significant role of cathepsin X in PD pathogenesis.

**Figure 8.**
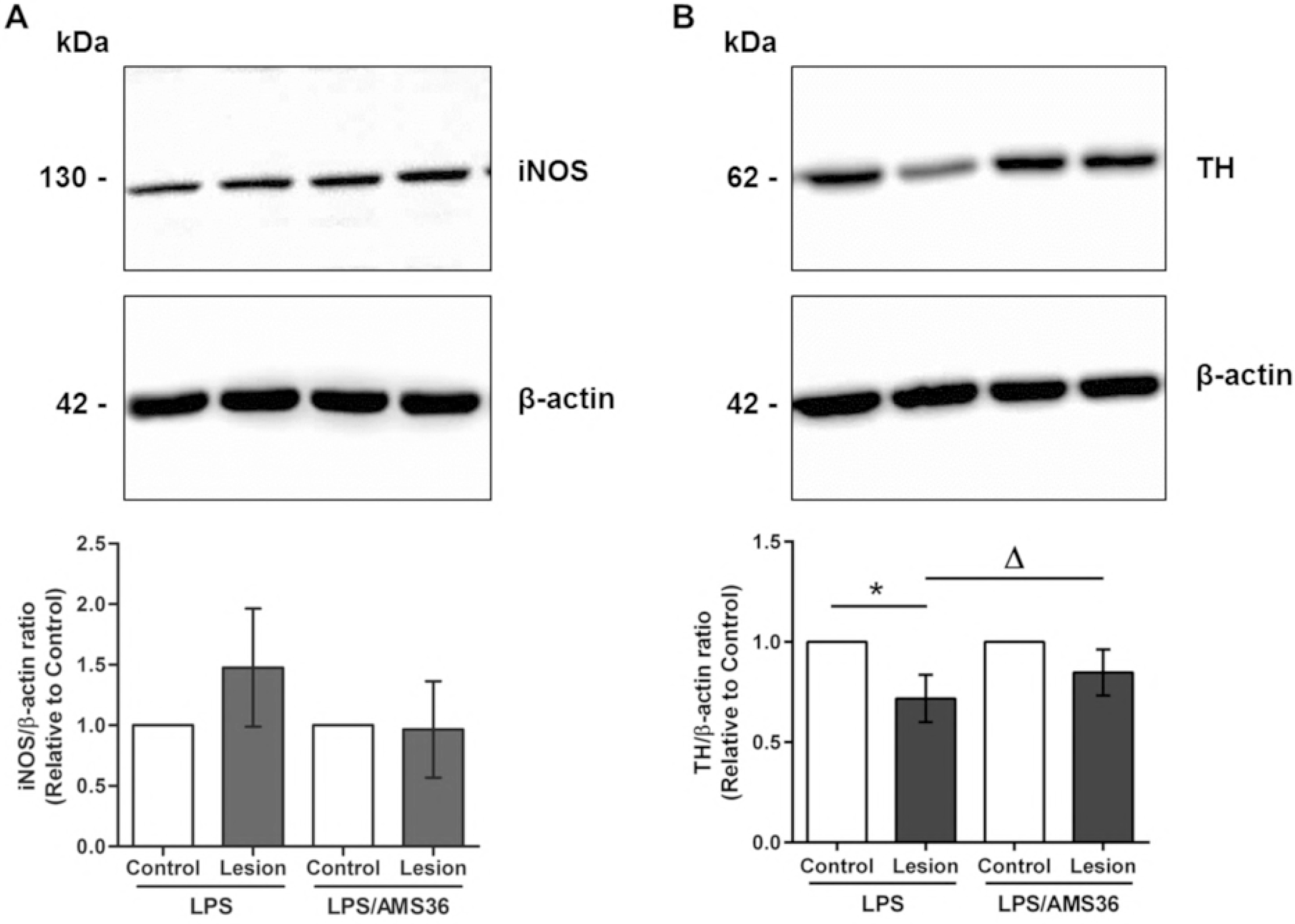
Cathepsin X inhibitor affected the protein levels of iNOS and TH in the ipsilateral striatum. (A, B) After LPS-induced lesion in the absence of cathepsin X inhibitor AMS36 (LPS) or the presence of AMS36 (LPS/AMS36), the contralateral (Control) and ipsilateral (Lesion) striata were dissected from coronal sections, homogenized, and analyzed for iNOS and TH protein levels. Western blot analysis was performed using rabbit polyclonal anti-iNOS antibody (A) and rabbit monoclonal anti-TH antibody (B). An antibody raised against β-actin was used as loading control. The graphs below blots show a semi-quantitative densitometry analysis of the protein level in the ipsilateral side relative to that in the contralateral side. Values are means ± SD of group of 5 animals for LPS and group of 6 animals for LPS/AMS36, where 4 anterior-posterior striatal slices each animal were analyzed (n = 5/6) (two-tailed Student’s *t*-test, * *p* < 0.05 *vs* Control; ^Δ^ *p* < 0.1 LPS/AMS36 *vs* LPS).

## Discussion

In this study, we utilized a model of neuroinflammation-induced degeneration of rat brain by intrastriatal injection of LPS, and aimed to evaluate the regional and cellular expression and activity of lysosome cysteine cathepsin X. We described for the first time that cathepsin X expression and its activity were strongly upregulated in the striatum after LPS-induced lesion. Along with striatal upregulation, cathepsin X overexpression was also found in the cortex, corpus callosum, subventricular zone and external globus pallidus, and in all of these regions the upregulation was restricted to glial cells. Moreover, we explored the biological role of cathepsin X in neuroinflammation by cathepsin X-specific inhibitor AMS36. The latter showed moderate protective effect toward the striatal lesion, which was reflected in reduced expression of the neuroinflammation marker iNOS. Therefore, we suggest that cathepsin X might play a significant role in neuroinflammation related to neurodegeneration.

Over the past decade, increasing evidence has emerged that implicates the role of cathepsin X in the central nervous system (CNS) neurodegeneration. High cathepsin X levels have been observed in degenerating brain regions of amyotrophic lateral sclerosis and AD transgenic mouse models and around senile plaques in brains of AD patients (Hafner et al, 2013; Wendt et al, 2007). Furthermore, a comprehensive comparative gene expression analysis of mouse models of AD, multiple sclerosis and stroke showed that cathepsin X is one of the eighteen genes whose expression is increased in all three models of CNS disorders (Tseveleki et al, 2010). Cathepsin X has also been associated with inflammation processes leading to neurodegeneration. It has been shown that cathepsin X is disproportionately expressed and secreted by microglia and astrocytes in response to neuronal damage and inflammatory stimulus, both in culture and *in vivo* (Glanzer et al, 2007; Greco et al, 2010; Pislar et al, 2017; Wendt et al, 2009). It was reported that dendritic cells in aging mice brains show increased expression of cathepsin X protein level that correlates with known markers of neuroinflammation (Stichel & Luebbert, 2007). Additionally, Allan et al. (Allan et al, 2017) showed that mice deficient in cathepsin X have reduced neuroinflammation and dramatically decreased circulating levels of interleukin 1β during experimental autoimmune encephalomyelitis. We recently showed the upregulation of cathepsin X in degenerated rat brain using a 6-OHDA rat PD experimental model. Unilateral 6-OHDA-induced lesion of the nigrostriatal pathways rapidly increased cathepsin X expression and activity in the ipsilateral SNc, which was mainly localized in TH-positive dopaminergic neurons, whereas a late time point of 6-OHDA-induced lesion caused persistent cathepsin X upregulation restricted to activated glia cells (Pislar et al, 2018). Thus, cathepsin X could be involved in neuroinflammation-induced dopaminergic neurodegeneration.

In this study, we adopted a neuroinflammation rat model of PD (Choi et al, 2009). Unilateral injection of LPS into the striatum leads to neuroinflammation-induced progressive dopaminergic neurodegeneration, which is accompanied by a progressively decreased number of TH-positive SNc cells, as well as by decreased TH-positive fibers in the striatum and decreased striatal DA levels (Choi et al, 2009; Hunter et al, 2009). In our study, intrastriatal LPS injection induced striatal degeneration as observed by reduced striatum region and increased size of the lateral ventricle four weeks after injection. In addition, a marked decrease of TH-positive fibers in the ipsilateral striatum was observed; however, there was no obvious loss of TH-positive cells in the ipsilateral SNc following intrastriatal LPS injection. The latter observation is consistent with the study of Hoban et al. (Hoban et al, 2013), who showed that intrastriatal LPS administration did not lead to any nigrostriatal neurodegeneration. However, rats with instrastriatal LPS injection rotated significantly after amphetamine administration, a behavioral indicator of induced functional alterations in the nigrostriatal dopaminergic system on the ipsilateral side. Thus, LPS treatment enhances responding to amphetamine after LPS treatment, suggesting that LPS administration dysregulates presynaptic DA levels, post-synaptic DA receptor sensitization or alterations in DA turnover (Fan et al, 2011; Fortier et al, 2004; Hoban et al, 2013).

Whilst the expression and release of cathepsin X has been shown to be associated with neuroinflammation (Glanzer et al, 2007; Greco et al, 2010; Pislar et al, 2017; Wendt et al, 2009), this study provides the first experimental evidence that cathepsin X is strongly upregulated in rat brain following LPS-induced striatal inflammation leading to neurodegeneration. We showed a marked increase in protein expression in the ipsilateral striatum and surrounding areas four weeks after LPS injection. Likewise, TH staining showed that there is probably no loss of dopaminergic cells in the ipsilateral SNc, as well as no change in cathepsin X immunoreactivity in the ipsilateral SNc four weeks after LPS injection. Additionally, we showed that cathepsin X protein level was significantly increased in the striatum on the lesioned side as compared to the contralateral striatum, where the pro-form of upregulated cathepsin X was most prominent. Moreover, our enzyme assays showed a significant increase of cathepsin X activity in the ipsilateral striatum following LPS injection, which implicates that cathepsin X might display a significant proteolytic activity in the analyzed brain region during neuroinflammation-induced neurodegeneration.

Alterations in the expression patterns and localization of the lysosomal cathepsins in the CNS have been reported in normal and aged brain, and under pathological conditions (reviewed in (Pislar & Kos, 2014)). Cathepsin X expression and proteolytic activity were detected in different brain regions, such as cerebral cortex, the cerebellum, the brainstem and the spinal cord in the adult mouse brain, and were found to be age-dependently upregulated during degenerative processes throughout brain regions (Wendt et al, 2007). Here, we examined the abundance and the regional distribution of cathepsin X in the lesioned striatum following LPS-induced neuroinflammation. Our immunohistochemical data show an abundant increase in cathepsin X expression predominantly in the ipsilateral striatum where condensed expression of cathepsin X was apparent or scattered throughout the striatum region. Nevertheless, strong upregulation of cathepsin X was also observed throughout different areas, such as in the cortex and corpus callosum around the injection track and in the external globus pallidus, where scattered distribution of cathepsin X through the region was observed. Thus, our data add another enzyme to the group of cysteine cathepsins, containing cathepsin B (Yan et al, 2013), L (Xu et al, 2018), H (Fan et al, 2015) and C (Fan et al, 2012), that have been reported to be upregulated in different brain regions following LPS-induced neuroinflammation.

Several lines of evidence support the hypothesis that glial reactive and inflammatory processes contribute in the cascade of events leading to neuronal degeneration (reviewed in (More et al, 2013)). The presence of glial cells, including reactive astrocytes, was observed in the brains of patients with PD (Cabezas et al, 2014; Teismann et al, 2003). Likewise, an increased number of activated microglial cells has consistently been reported in the neuroinflammatory pathogenesis of PD (Banati et al, 1998; Choi et al, 2009). Microglia, which are the most abundant glia cells in the brain, have been observed to grow densely in the striatum and SNc with increased expression of proinflammatory mediators (Hoban et al, 2013; Tansey et al, 2007). LPS injection into the striatum caused striatal microgliosis, and the presence of reactive astrocytes and we found that the majority of cathepsin X-immunoreactive signals were localized in these glial cells in the area of the damaged striatum. Cathepsin X staining was localized mainly within microglia cells in the striatum and throughout different areas including the cortex, corpus callosum, subventricular zone and external globus pallidus. This observation correlates well with previous *in vitro* studies showing upregulated expression, increased release and activity of microglial cathepsin X following LPS stimulation (Pislar et al, 2017; Wendt et al, 2009). We also observed cathepsin X upregulation in reactive astrocytes, where stronger co-localization with astrocytes marker was observed in the subventricular zone; however, some astrocytes in the ipsilateral striatum and surrounding areas also showed cathepsin X overexpression. On the contrary, no neuronal cells were positive for upregulated cathepsin X at the ipsilateral striatum and throughout other regions following LPS injection. These results are in agreement with our recent *in vivo* study using a hemi-parkinsonian 6-OHDA rat model of PD. Four weeks post-lesion, we observed a persistent cathepsin X upregulation restricted to glial cells concentrated in the ipsilateral SNc (Pislar et al, 2018). The data obtained in the present study are in line also with other reports showing glial upregulation of certain cysteine cathepsins in LPS-induced neuroinflammation (Fan et al, 2015; Fan et al, 2012).

Due to the harmful action of cysteine cathepsins in pathological processes of neurodegeneration, cathepsin inhibitors constitute a possible tool for therapeutic interventions to inhibit excessive proteolytic activity (reviewed in (Pislar & Kos, 2014). Some beneficial *in vivo* effects of cysteine protease inhibitors towards neurodegeneration have been demonstrated with cathepsin B inhibition (Haque et al, 2008; Hook et al, 2007; Hook et al, 2010; Van Broeck et al, 2007). The results obtained so far implicate a significant role of cathepsin X in neuroinflammation-induced neurodegeneration; therefore, the inhibition of its excessive activity might be a useful therapeutic approach. Here, we used a known irreversible epoxysuccinyl-based inhibitor of cathepsin X AMS36 (Sadaghiani et al, 2007), which demonstrated promising neuroprotective effects *in vitro* (Pislar et al, 2017; Pislar et al, 2014). We demonstrated lower cathepsin X activity in rat brain after AMS36 administration as well as selectivity for cathepsin X among other cysteine cathepsins. AMS36 selectivity towards cathepsin X was also reported for tumor tissues, whereas in other tissues such as rat liver and kidney, a significant cross-reactivity of AMS36 with cathepsin B inhibition was observed (Sadaghiani et al, 2007). Nevertheless, AMS36 exerted a moderate protective effect on LPS-induced striatal degeneration based on the observed trend toward the reduced size of the lateral ventricle and decreased extent striatal lesion. Additionally, continuous administration of AMS36 along with LPS injection also affected the protein levels of TH and iNOS. To date, one of the best elucidated cytotoxic mechanisms induced by proinflammatory cytokines in neuroinflammation related to PD is activation of iNOS. The protein level of iNOS is increased in both the SNc and striatum after LPS challenge (Choi et al, 2009; Dawson et al, 1993). Inhibition of cathepsin X by AMS36 reversed the increased iNOS protein level in the ipsilateral striatum, which coincided with increased TH protein level in the ipsilateral striatum following LPS injection in presence of AMS36. Although the evaluated cathepsin X inhibitor did not exhibit significant protective effects towards striatal LPS-induced neuroinflammation, the results clearly identify cathepsin X as a key player in neuroinflammation-induced neurodegeneration. A new generation of selective and reversible cathepsin X inhibitors (Fonovic et al, 2017) may be expected to significantly improve the cathepsin X-targeted therapy of neurodegenerative diseases such as PD.

Taken together, in our current study, we established an animal model of neuroinflammation by intrastriatal LPS administration. The findings show the regional distribution and cellular localization of cathepsin X, a lysosomal cysteine protease, in the brain. LPS injection into the striatum increased cathepsin X expression and its activity in the striatum and surrounding areas on the ipsilateral side. The prominent cathepsin X upregulation was restricted to glial cells, activated microglia and reactive astrocytes. Moreover, administration of a cathepsin X inhibitor along with LPS injection exerted a trend towards decreased extent of striatal lesion and decreased expression of neuroinflammation marker iNOS. These results add to previous *in vitro* evidence showing cathepsin X mediated microglia activation and further support the notion that cathepsin X could participate in the development of neuroinflammation in the CNS.

## Materials and Methods

### Animals

Male Wistar rats, from the Medical Experimental Center, Medical Faculty, University of Ljubljana, Slovenia, weighing 200-250 g at the beginning of the experiments were housed at 22 ± 2 °C and relative humidity 60 ± 10% under a 12 h light/dark cycle, with free access to food and water. All animal-related procedures were conducted in accordance with the European Communities Council Directive of 2010 (2010/63/UE) and the National Veterinary Institute Guide for the Care and Use of Laboratory Animals. Care was taken to minimize the number of experimental animals and their suffering. The experiments were approved by the Administration of the Republic of Slovenia for Food Safety, Veterinary Sector and Plant Protection.

### Intrastriatal lipopolysaccharide-induced lesion

Surgical procedures for intrastriatal LPS injection were adopted from Choi et al. (Choi et al, 2009). For the injection of LPS (Salmonella Minnesota; Sigma Aldrich, St. Louis, MO, USA), male Wistar rats (n=5-6/group) were deeply anesthetized with sodium phenobarbital (Sigma-Aldrich; 50 mg/kg i.p.), and then positioned in a stereotaxic frame (TrentWells, South Gate, CA, USA) with the incisor bar at the level of the ear. LPS dissolved in saline (2.5 μg/μL) was injected into the right striatum (3 μL/site) at the following coordinates (in mm): site 1, anteroposterior (AP) 1.0, mediolateral (ML) 2.0, dorsoventral (DV) −4.5; site 2, AP 1.0, ML 3.0, DV −5.5; site 3, AP −0.5, ML 3.0, DV −4.5; site 4, AP −0.5, ML 4.0, DV −6.0. Saline was injected into the left striatum with parallel coordinates. The cannula was left at the injection site for an additional 3 min post-injection before being slowly retracted. After surgery, animals were kept on a heating pad until recovery from surgery and subcutaneous saline was given for aid in postoperative recovery.

In order to investigate the effect of cathepsin X inhibition on the LPS-induced lesion, a specific, irreversible inhibitor of cathepsin X AMS36, synthesized as reported (Pislar et al, 2014), was used. One group of male Wistar rats (n=6) was treated every fourth day for 4 weeks with the inhibitor AMS36 in dimethyl sulfoxide (DMSO)/saline solution ratio 1:10 (vol/vol) at a dose of 50 mg/kg (i.p.). For vehicle control, the other group of male Wistar rats (n=5) received DMSO/saline solution in the same manner. The first injection of AMS36 or vehicle was performed 2 days before LPS injection, and the last 2 days prior to animal sacrifice.

### Brain tissue preparation

Subjects were sacrificed after 4 weeks of LPS injection and brains were rapidly removed and quickly frozen on dry ice and stored at −80°C until sectioning on a cryostat. Coronal sections (10 – 20 μm), identified using a rat brain atlas (Paxinos & Watson, 2007), were cut at four anterior-posterior levels through the striatum (in mm from bregma); 1) between 1.92 and 1.20, 2) between 1.08 and 0.24, 3) between 0.12 and −0.48 and 4) between −0.60 and −1.44 and at three anterior-posterior levels through the SNc (in mm from bregma); 1) between 4.68 and −5.04, 2) between −5.20 and −5.52 and 3) between −5.64 and −6.00. The sections were mounted onto microscope glass slides coated with a 0.01% solution of (poly)L-lysine (Sigma-Aldrich) or adhered to parafilm. The slides were then vacuum-packed and stored in a freezer at −20 °C until being further processed.

### Methylene staining

Slices (10 μm) were fixed in 4% formaldehyde in 0.1 M sodium phosphate buffer saline (NaPBS; pH 7.2– 7.4, 4 °C) for 5 min. After washing with potassium phosphate buffer saline (KPBS, pH 7.2), sections were counterstained with 0.2% methylene blue (Sigma-Aldrich) for 1 min, rinsed with distilled water, dehydrated in rising ethanol concentration and coverslipped with the DPX mounting medium (BDH Laboratory Supplies, Poole, UK).

### Lateral ventricle and striatum area quantification

Methylene blue-stained brain slices were scanned by MCID, M4 analyzer (Imaging Research, St Catharines, ON, Canada). Areas of lateral ventricles and striata were manually outlined and measured in striatal brain slices cut at 4 different anterior-posterior levels. The scan area (pixels) of lateral ventricle/striatum was separately measured in the contralateral and ipsilateral side. In the graphs sum of scan area of lateral ventricles/striata in striatal brain slices are shown.

### Immunohistochemistry

Coronal brain sections (10 μm) were fixed in cold 100% methanol for 5 min followed by 15 min in cold methanol with 1% H_2_O_2_. Brain slices were incubated in blocking buffer containing 4% normal serum, 1% bovine serum albumin (BSA), and 0.1% Triton X-100 in KPBS for 1 h at room temperature. They were then incubated overnight at 4°C with mouse monoclonal antibody against tyrosine hydroxylase (TH) (1:750, Abcam, Cambridge, UK) or goat polyclonal primary antibody against cathepsin X (1:200, AF934, R&D Systems, MN, USA), diluted in blocking solution. Afterwards, the sections were incubated with biotinylated anti-mouse or anti-goat secondary antibodies (1:750, Vector Laboratories, Burlingame, CA, USA) diluted in KPBS containing 1% BSA and 0.02% Triton X-100 for 1.5 h at room temperature. Avidin-biotin-peroxidase complex (ABC elite standard kit, Vector Laboratories, Burlingame, CA, USA) was added for 30 min. Staining was visualized with 3,3′-diamino-benzidine (DAB, Sigma-Aldrich). All sections were simultaneously immunolabeled to ensure identical DAB staining incubation times. Sections were then dehydrated and coverslipped with DPX mounting medium (BDH Laboratory Supplies). Brain sections were examined and imaged with an Olympus microscope (Olympus IX81) with an attached digital camera (Olympus DP71) using the same system settings for all samples.

### Protein extraction from striatal and nigral sections

For analysis of the protein level of cathepsin X and its activity, the striatum and SNc were dissected out from four 20 μm frozen brain slices (striatum) or eight 20 μm frozen brain slices (SNc) of LPS or LPS/AMS36 rat brains, separately from the contralateral (Control) and ipsilateral side (Lesion), using a cryostat at −20 °C. Tissue was homogenized in ice-cold lysis buffer (0.05 M sodium acetate, pH 5.5, 1 mM EDTA, 0.1 M NaCl, 0.25% Triton X-100) supplemented with a cocktail of phosphatase inhibitors (Thermo Fisher Scientific, Waltham, MA, USA), then sonicated and centrifuged at 15.000 *g* at 4 °C for 15 min to collect the supernatant. Total protein concentration was determined by DC™ Protein Assay (Bio-Rad, Hercules, CA, USA). All the samples were kept at −70°C until they were used for analysis.

### Western blotting

The protein levels of TH, cathepsin X and inducible nitric oxidase synthase (iNOS) were determined by western blotting as previously reported (Hafner et al, 2012). Separated proteins, transferred to a nitrocellulose membrane were immunoblotted with the rabbit monoclonal anti-TH antibody (1:400, Abcam), goat polyclonal anti-cathepsin X antibody (1:500, R&D Systems), rabbit polyclonal anti-iNOS (1:500, Abcam) and mouse monoclonal anti-β-actin antibody (1:500, Sigma Aldrich). Signals from anti-goat conjugated with horseradish peroxidase (HRP) (1:3000, Santa Cruz Technology), anti-rabbit conjugated with HRP or anti-mouse conjugated with HRP (1:5000, Millipore, Billerica, MA, USA) secondary antibodies were visualized with an enhanced chemiluminescence detection kit (Thermo Scientific, Rockford, IL, USA). The band intensities were quantified using Gene Tools software (Sygene, UK), and expressed as values relative to those of controls.

### ELISA

The protein level of cathepsin X was determined by ELISA as previously reported (Kos et al, 2005). Briefly, microtiter plates were coated with equal aliquots of goat polyclonal anti-cathepsin X antibody (RD Systems) in 0.01 M carbonate/bicarbonate buffer, pH 9.6, at 4 °C. After blocking with 2% BSA in phosphate buffered saline (PBS), pH 7.4, for 1 h at room temperature, the samples of equal protein amount (50 μg) or cathepsin X standards (0 – 65 ng/mL) were added. Following 2 h incubation at 37 °C, the wells were washed and filled with mouse monoclonal anti-cathepsin X 3B10 antibody conjugated with HRP in blocking buffer. After a further 2 h incubation at 37 °C, 200 μL/well of 3,3′,5,5′-tetramethylbenzidine (TMB) substrate (Sigma-Aldrich) in 0.012% H_2_O_2_ was added. After 15 min, the reaction was stopped by adding 50 μL/well of 2 μM H_2_SO_4_. The amount of protein was determined by measuring the absorbance at 450 nm using a microplate reader (Tecan Safire^2^, Switzerland), and the concentration of cathepsin X was calculated from the standard calibration curve.

### Cathepsin X activity

Cathepsin X activity was measured in tissue lysates with the cathepsin X-specific, intramolecularly quenched fluorogenic substrate Abz-Phe-Glu-Lys(Dnp)-OH synthesized by Jiangsu Vcare Pharmatech Co. (China). An aliquot of 50 μg of the lysate proteins was incubated at 37 °C, followed by measurement of cathepsin X activity using 10 μM Abz-Phe-Glu-Lys(Dnp)-OH. The fluorometric reaction was quantified at 37 °C at an excitation wavelength of 320 nm and emission wavelength of 420 nm on a microplate reader (Tecan Safire^2^). Results are presented as change in fluorescence as a function of time (ΔF/Δt) and cathepsin X activity was expressed relative to Control.

### Double immunofluorescence labeling

The striatal and nigral sections (10 μm) of LPS treated rats were double-immunostained for the analysis of the cellular localization of cathepsin X using the following primary antibodies: goat polyclonal anti-cathepsin X antibody (1:75, R&D System), mouse monoclonal anti-TH antibody (1:500, Abcam) as marker for dopaminergic neurons, mouse monoclonal anti-NeuN antibody (1:300, EMD Millipore) as neuronal marker, mouse monoclonal anti-CD11b antibody (1: 175, Abcam) as microglia marker and mouse monoclonal anti-glial fibrillary acidic protein (GFAP) antibody (1:1000, Abcam) as astrocyte marker. Brain slices were fixed in cold methanol for 20 min and the immunofluorescence procedure was further performed as reported (Tratnjek et al, 2016). Briefly, brain slices were then incubated in blocking solution containing 4% donkey serum (EMD Millipore) and 0.4% Triton X-100 in KPBS for 1 h at room temperature. Brain sections were then incubated with the primary antibodies overnight at 4°C diluted in KPBS containing 1% donkey serum and 0.4% Triton X-100, followed by 1.5 h incubation at room temperature. Thereafter, sections were incubated with the Alexa-fluorophore-conjugated secondary antibodies (1: 300, Invitrogen, Molecular Probes, OR, USA) diluted in KPBS containing 0.02% Triton X-100 for 1.5 h at room temperature. After the incubation, sections were immersed into 0.1% Sudan Black B (Sigma-Aldrich) in 70% ethanol (vol/vol) for 5 min to suppress lipofuscein autofluorescence background.

Sections were then rinsed with KPBS and coverslipped using the ProLong® Gold Antifade Mountant with DAPI (Thermo Fisher Scientific). To confirm staining specificity, primary antibodies were omitted in the staining procedure (*data not shown*). Fluorescence microscopy was performed using a Carl Zeiss LSM 710 confocal microscope (Carl Zeiss, Oberkochen, Germany). Images are presented as single confocal sections and were analyzed using Carl Zeiss ZEN 2011 image software. All images were captured under the same exposure time settings. To improve the signal/noise ratio, 4 frames/image were averaged.

### Statistical analysis

Results of protein levels and cathepsin X activity are representative of two independent experiments, each performed in duplicate, and are presented as means ± SD. Statistical evaluation was performed with the two-tailed Student’s *t*-test when two values sets were compared, where *p* < 0.05 was considered to be statistically significant. The differences in striatum area or size of lateral ventricles between contralateral and ipsilateral sides within each experimental group (LPS, LPS/AMS36) were analyzed using ANOVA test and post hoc analysis using Bonferroni multiple comparisons *t*-test. When two sets of values were compared; *p* < 0.05 was considered to be statistically significant. For all the statistical evaluation, GraphPad Prism (version 6) was used.

## Acknowledgements

We acknowledge Prof. Ronald E. See for critical review of the paper before submission. This study has been supported by the Research Agency of Republic of Slovenia (Grants P4-0127 and J4-4123 to JK and P3-0171 to RS).

## Author contributions

AP, MŽ and JK designed the study. AP prepared coronal sections and protein extractions, performed immunohistochemistry staining, Western blotting, ELISA assay, cathepsin X activity assay and double immunofluorescence labeling, generated the data for Figures and prepared the draft manuscript. LT performed methylene staining and lateral ventricle and striatum area quantification, immunohistochemistry analysis, participated in carrying out the double immunofluorescence analysis and reviewed the manuscript. GG reviewed the manuscript. NZ synthesized the cathepsin X inhibitor AMS36. MŽ performed animal surgery. MŽ and JK coordinated the research and reviewed the manuscript. All authors read and approved the final manuscript.

## Conflict of interest

The authors declare the absence of competing interest.

